# *In silico* identification of miRNAs related to mitochondrial dysfunction in amyotrophic lateral sclerosis

**DOI:** 10.1101/2022.09.05.506596

**Authors:** Baykal Gulcin, Erkal Burcin, Vural Korkut Senay

## Abstract

Non-coding, single-stranded RNA molecules known as microRNAs (miRNAs) regulate gene expression via mRNA degradation after transcription. As a result, they affect a number of pathways in organisms that are important for both health and disease. miRNAs can be utilized as potential diagnostic, prognostic, and therapeutic biomarkers for neurodegenerative diseases such as Amyotrophic Lateral Sclerosis (ALS). Neuronal cells are highly dependent on mitochondria, and mitochondrial dysfunction has been linked to neurodegenerative diseases. Pathological changes in ALS are associated with disruptions in mitochondrial structure, bioenergetics, and calcium homeostasis. In this study, we used an *in silico* approach to identify miRNAs associated with mitochondrial dysfunction in ALS based on target genes that are implied in both ALS and mitochondrial dysfunction. A literature search revealed the genes SOD1, FUS, TARDBP, C9orf72, CHCHD10, OPTN, VCP, TBK1 and BCL2 that cause mitochondrial dysfunction and are involved in the pathogenesis of ALS. Pathway enrichment analyses using Enrichr, g:Profiler, and CROssBAR tools confirmed that the identified genes have significant associations with ALS, mitochondrial dysfunction, and neuron differentiation. In silico miRNA predictions have been made using the databases miRWalk, miRTargetLink, TargetScan, and miRNet. A Venn diagram tool was used to select common miRNAs, and finally 28 miRNAs were discovered. One set of 28 miRNAs were subjected to set analysis using the miRNet and TAM tools for functional and enrichment analyses, respectively. In both databases, three common miRNAs, hsa-miR-9-5p, hsa-miR-141-3p and hsa-miR-125b, were found to be linked to ALS.

## 1. Introduction

Amyotrophic Lateral Sclerosis (ALS) is a fatal motor neuron disease that results from the degeneration of upper and lower motor neurons. ALS is characterized by progressive muscular atrophy affecting motor neurons in the cerebral cortex, brainstem, and spinal cord, resulting in muscle twitching and neurons losing their synaptic connections with the target muscles. This disease usually manifests itself in late middle age (Brown and Al-Chalabi, 2017; Hardiman et al., 2017; Mejzini et al., 2019). While more than 90% of ALS patients are sporadic (sALS), 5– 10% of the total population of ALS patients may have a familial form of the disease. According to epidemiological studies conducted in many countries, the incidence of ALS ranges from 0.6 to 3.8 per 100,000 people, and the prevalence ranges from 4.1 to 8.4 per 100,000 people (Benjaminsen et al., 2018; Jun et al., 2019; Leighton et al., 2019; Longinetti et al., 2018; Palese et al., 2019; Rose et al., 2019; Zhou et al., 2018). Both the incidence and prevalence of the disease increase in parallel with age (Brown and Al-Chalabi, 2017).

Many cellular and molecular processes are involved in the pathophysiology of ALS, including mitochondrial dysfunction, axonal transport, toxic protein aggregation, excitotoxicity, oxidative stress, RNA metabolism defects, and inflammation (van Es et al., 2017; Hardiman et al., 2017). Neurons are cells with many mitochondria due to their high energy and oxygen requirements, and as a result, they are more vulnerable to oxidative damage (Bordoni and Gabbianelli, 2020; Lezi and Swerdlow, 2012). Mitochondrial dysfunction is linked to a variety of pathological conditions, including cancer, cardiovascular disease, and diabetes, and is crucial in the development of various neurodegenerative diseases (Morgan and Orrell, 2016; Peoples et al., 2019; Wallace, 2012). These dysfunctions not only disrupt the mitochondrial energy mechanism, but also cause problems with mitophagy, protein processing in endoplasmic reticulum (ER), and calcium signalling (Jaiswal, 2017; Morgan and Orrell, 2016; Tadic et al., 2014).

As early as 1981, mitochondria with abnormal size and shape were discovered in intramuscular nerves from 11 ALS patients’ muscle biopsies (Atsumi, 1981). An ultrastructural examination of the synapses on the surface of the somata of anterior horn neurons in the lumbar spinal cord of seven ALS patients was later performed, and dense conglomerates of aggregated dark mitochondria were discovered (Sasaki and Iwata, 1996). More detailed studies on the mitochondrial changes in nerve tissues obtained from ALS patients were conducted in subsequent years. In such a study, the anterior horns of the lumbar spinal cord were examined by electron microscopy in 14 patients with sporadic ALS, and abnormal alterations were found in half of the patients. These included changes in the intermembrane space and the inner mitochondrial membrane (Sasaki and Iwata, 2007). Lin’s study was the first to obtain blood samples, and it discovered ETS dysfunction as a result of decreases in the enzyme/protein levels of oxidative phosphorylation in blood samples taken from ALS patients (Lin et al., 2009).

MicroRNAs (miRNAs) are single-stranded, non-coding RNA molecules that regulate gene expression post-transcriptionally. A single miRNA can regulate many target mRNAs, and an mRNA transcript can be regulated by many miRNAs (Bartel, 2004; Peter, 2010; Wu et al., 2010). miRNAs bind to the 3’ untranslated region (UTR) of the target mRNA, showing complete or partial complementarity. It is critical that miRNAs with a seed sequence of 2-8 nucleotides complement the target mRNA (Bartel, 2009; Grimson et al., 2007). Some miRNAs are involved in neurogenesis (Åkerblom et al., 2012) and neuronal differentiation (Conaco et al., 2006).

Early ALS diagnosis, which is critical for initiation of treatment, is a long process and takes an average of 12 months (Brown and Al-Chalabi, 2017). In this case, biomarker research has priority in ALS studies (Bowser et al., 2011; van Es et al., 2017).

Due to the complexity of gene expression in ALS-related pathways, there are currently no validated conventional biomarkers for the diagnosis or monitoring of ALS disease progression. In this study, we used *in silico* analyses to identify specific miRNAs associated with mitochondrial dysfunction in ALS, based on miRNA target genes that are important in both ALS and mitochondrial dysfunction. Our aim is to show that common miRNAs identified by bioinformatics tools can be utilised as potential biomarkers to shorten and simplify the diagnosis of ALS disease.

## 2. Materials and Methods

### Data Collection and Literature Search Strategy

A literature search on mitochondrial dysfunction and associated genes in ALS disease was conducted through electronic searches in PubMed and Science Direct databases for eligible studies published up to 2022. We used the MeSH terms that are ALS, microRNAs, and mitochondria.

### Pathway Enrichment Analyses of Determined Genes

The determined genes from the literature search were run through the Enrichr (https://maayanlab.cloud/Enrichr/) gene set enrichment analysis tool. Pathway analysis in KEGG 2021 Human was performed on the identified genes. The determined genes were also run in the g:Profiler (https://biit.cs.ut.ee/gprofiler/gost) tool. Biological pathway analyses were carried out using KEGG and WikiPathway in g:Gost functional profiling. Finally, the genes were run through the CROssBAR (https://crossbar.kansil.org) tool.

### miRNA Prediction and Eligibility Criteria

To obtain more specific results, four different databases, miRWalk (http://mirwalk.umm.uni-heidelberg.de), miRTargetLink 2.0 (https://ccb-compute.cs.uni-saarland.de/mirtargetlink2), miRNet 2.0 (https://www.mirnet.ca), and TargetScan (http://www.targetscan.org/vert_80/), were used to identify miRNAs targeting the determined genes. All databases have been searched for *homo sapiens*. Each identified gene was run through four databases separately. For each gene, a Venn diagram tool (https://bioinformatics.psb.ugent.be/webtools/Venn/) was used to visualize common miRNAs related to the corresponding gene using miRNAs obtained from four separate databases. This research was carried out within certain inclusion criteria: i) miRNAs found in at least two databases for each gene were collected into a single list, ii) miRNAs from the common list that targeted at least three genes were combined into a single list (Table S1). For all process *p<0*.*05* was accepted statistically significant.

### miRNA Set Analyses

Following the creation of the common list, set analysis was carried out using the miRNAs by the miRNet 2.0 (https://www.mirnet.ca) and TAM 2.0 (http://www.lirmed.com/tam2/) tools shown in Table 3. As a result of previous enrichment analysis, relevant miRNAs in tissue, function, and disease areas were discovered in both databases, and a venn diagram was used to illustrate common miRNAs.

## 3. Results

### Selection of Genes

A literature search was conducted on genes associated to mitochondrial dysfunction in ALS. As a result of these studies the *SOD1, FUS, TARDBP, C9orf72, OPTN, TBK1, CHCHD10, VCP*, and *BCL2* genes were revealed, and research continued these genes (Table 1). Identified genes were labelled in yellow in the KEGG Pathways diagram (Figure 2).

**Table 1:** Genes are Related to Mitochondrial Dysfunction in ALS

| Locus | Gene (Protein) | Refs |
| --- | --- | --- |
| ALS1 | SOD1 (Superoxide dismutase 1) | Kong & Xu, 1998; Vande Velde et al., 2011 |
| ALS6 | FUS (RNA-binding protein FUS) | Sharma et al., 2016 |
| ALS10 | TARDBP (TAR DNA-binding protein 43) | W. Wang et al., 2016; Zuo et al., 2021 |
| ALS12 | OPTN (optineurin) | Wong & Holzbaur, 2014; Liu et al., 2018 |
| FTD/ALS 6 | VCP (valosin-containing protein) | Johnson et al., 2010; N. C. Kim et al., 2013 |
| FTD/ALS 1 | C9orf72 (guanine nucleotide exchange C9orf72) | Dafinca et al., 2016 |
| FTD/ALS 2 | CHCHD10 (coiled-coil-helix-coiled-coil-helix domain-containing 10) | Bannwarth et al., 2014; Genin et al., 2019 |
| FTD/ALS 4 | TBK1 (serine/threonine-protein kinase TBK1) | Harding et al., 2021 |
| 18q21.33 | BCL2 (B-cell Lymphoma) | Mu et al., 1996; Pedrini et al., 2010 |

### Pathway Enrichment Analyses of Determined Genes

The determined genes were carried out through KEGG analysis in the Enrichr database. The eight genes were found to be significantly associated with ALS. The seven genes were found to be strongly linked to neurodegeneration. The genes TBK1 and OPTN were found to be significantly associated with mitophagy, while TBK1 and BCL2 were found to be significantly associated with autophagy (Table 2).

**Table 2:** Enrichment pathway analysis for KEGG 2021 using the Enrichr tool.

| term | p value | q value | overlap genes |
| --- | --- | --- | --- |
| Amyotrophic Lateral Sclerosis | 9.877965e- 14 | 6.025558e- 12 | [VCP, TBK1, FUS, BCL2, CHCHD10, TARDBP, OPTN, SOD1] |
| Pathways of Neurodegeneration | 1.409405e- 10 | 4.298684e- 09 | [VCP, TBK1, FUS, BCL2, TARDBP, OPTN, SOD1] |
| Mitophagy | 4.037790e- 04 | 8.210173e- 03 | [TBK1, OPTN] |
| mRNA Surveillance Pathway | 8.365989e- 04 | 1.275813e- 02 | [FUS, TARDBP] |
| Autophagy | 1.624891e- 03 | 1.699943e- 02 | [TBK1, BCL2] |
| Measles | 1.672075e- 03 | 1.699943e- 02 | [TBK1, BCL2] |
| Hepatitis B | 2.261359e- 03 | 1.809705e- 02 | [TBK1, BCL2] |
| Protein Processing in Endoplasmic Reticulum | 2.515130e- 03 | 1.809705e- 02 | [VCP, BCL2] |
| NOD-like Receptor Signalling Pathway | 2.812236e- 03 | 1.809705e- 02 | [TBK1, BCL2] |
| RNA Transport | 2.966730e- 03 | 1.809705e- 02 | [FUS, TARDBP] |

The KEGG pathway analysis was performed on nine genes with g:Profiler tool. The nine genes appeared to be significantly associated with ALS. The eight genes were found to be strongly linked with neurodegeneration. TARDBP, TBK1, VCP, CHCHD10, C9ORF72, and SOD1 genes appeared to significantly linked with abnormality in mitochondrial morphology. As a result of this database analysis, it was revealed that there are many genes associated with the regulation of autophagy, neuron atrophy, and morphological abnormalities (Figure 3 new).

In addition, the CROssBAR tool was used for these genes. As a result, the TBK1 and OPTN genes were prominent in the mitophagy-related pathway. At the same time, TBK1, BCL2, and C9orf72 genes were found in autophagy-related pathway. Moreover, the BCL2 gene was also linked to the p53 signalling pathway and apoptosis. When BCL2 and VCP genes are evaluated together these genes were also found in protein processing in the endoplasmic reticulum (Figure 4). The KEGG pathway in figure 1 shows the p53 signalling pathway and protein processing in the endoplasmic reticulum in detail.

**Figure 1:**
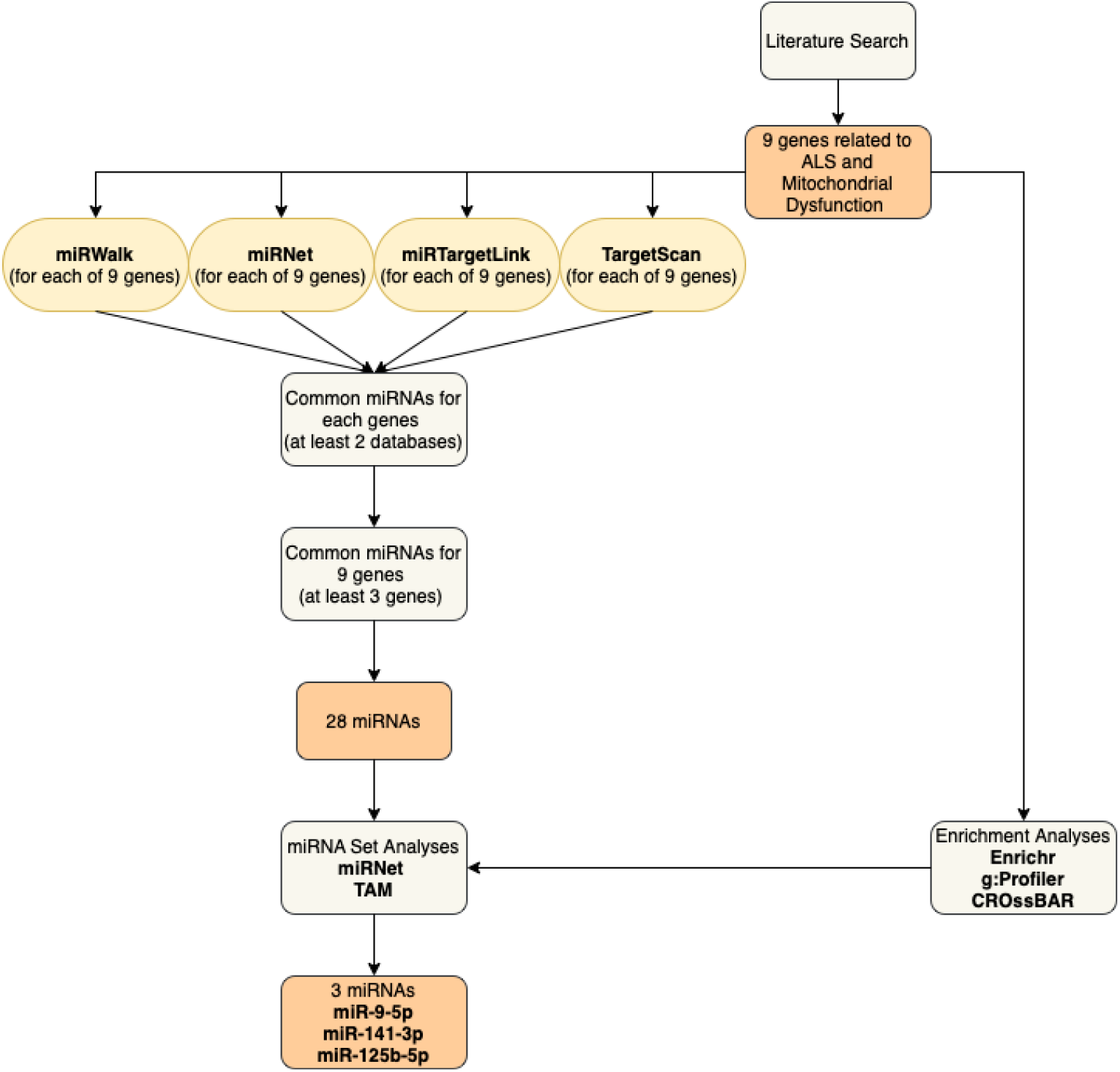
Overview of the study design

### Identification of miRNAs and Set Analyses

At the beginning, instead of a single database, four different databases were used to obtain more specific results.

Data was obtained from four databases separately for each gene. For each gene, a Venn diagram was used individually to find miRNAs that were found in at least two databases (Figure 5). All related miRNAs targeting at least three genes among common miRNAs found in at least two databases for each gene were listed using Venn diagram, as a result of analyses, 28 miRNAs were revealed (Table S1).

These miRNAs were also evaluated using the miRNet 2.0 and TAM 2.0 tools for miRNA set analysis. At this stage, miRNAs were eliminated using data from pathway enrichment analyses. When we compared the pathways associated with the nine identified genes and the pathways associated with the 28 miRNAs, we obtained the data in Table 3. Looking at the data in Table 3, we determined that miR-9, miR-141 and miR-125 were prominent. As a result, we obtained three miRNAs. Three common miRNAs, hsa-miR-9-5p, hsa-miR-141-3p and hsa-miR-125b-5p, were found to be significantly linked to ALS in both databases.

**Table 3:**
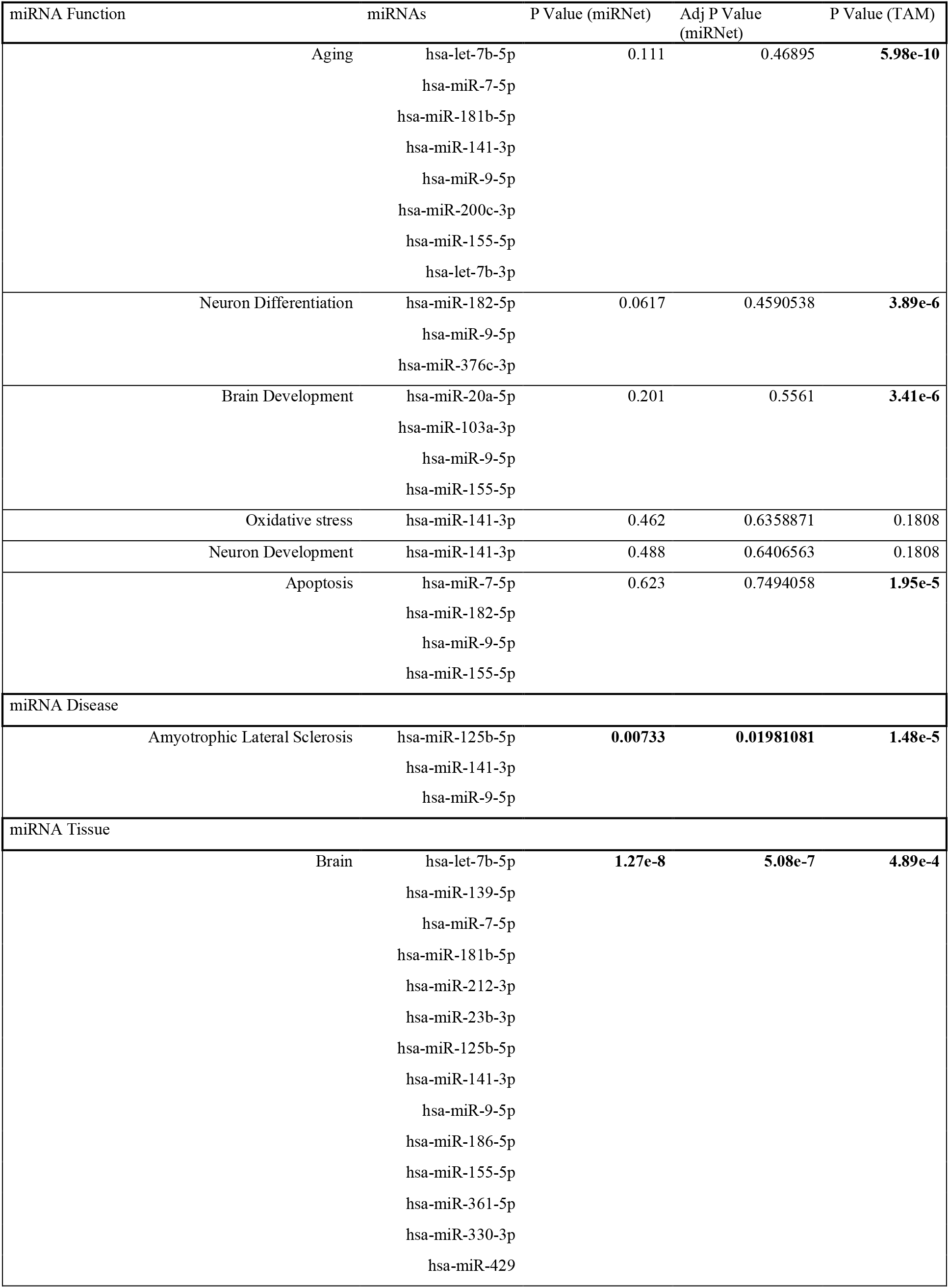
miRNet and TAM Set Analyses

| miRNA Function | miRNAs | P Value (miRNet) | Adj P Value (miRNet) | P Value (TAM) |
| --- | --- | --- | --- | --- |
| Aging | hsa-let-7b-5p | 0.111 | 0.46895 | **5.98e-10** |
|  | hsa-miR-7-5p |  |  |  |
|  | hsa-miR-181b-5p |  |  |  |
|  | hsa-miR-141-3p |  |  |  |
|  | hsa-miR-9-5p |  |  |  |
|  | hsa-miR-200c-3p |  |  |  |
|  | hsa-miR-155-5p |  |  |  |
|  | hsa-let-7b-3p |  |  |  |
| Neuron Differentiation | hsa-miR-182-5p | 0.0617 | 0.4590538 | **3.89e-6** |
|  | hsa-miR-9-5p |  |  |  |
|  | hsa-miR-376c-3p |  |  |  |
| Brain Development | hsa-miR-20a-5p | 0.201 | 0.5561 | **3.41e-6** |
|  | hsa-miR-103a-3p |  |  |  |
|  | hsa-miR-9-5p |  |  |  |
|  | hsa-miR-155-5p |  |  |  |
| Oxidative stress | hsa-miR-141-3p | 0.462 | 0.6358871 | 0.1808 |
| Neuron Development | hsa-miR-141-3p | 0.488 | 0.6406563 | 0.1808 |
| Apoptosis | hsa-miR-7-5p | 0.623 | 0.7494058 | **1.95e-5** |
|  | hsa-miR-182-5p |  |  |  |
|  | hsa-miR-9-5p |  |  |  |
|  | hsa-miR-155-5p |  |  |  |
| miRNA Disease |  |  |  |  |
| Amyotrophic Lateral Sclerosis | hsa-miR-125b-5p | **0.00733** | **0.01981081** | **1.48e-5** |
|  | hsa-miR-141-3p |  |  |  |
|  | hsa-miR-9-5p |  |  |  |
| miRNA Tissue |  |  |  |  |
| Brain | hsa-let-7b-5p | **1.27e-8** | **5.08e-7** | **4.89e-4** |
|  | hsa-miR-139-5p |  |  |  |
|  | hsa-miR-7-5p |  |  |  |
|  | hsa-miR-181b-5p |  |  |  |
|  | hsa-miR-212-3p |  |  |  |
|  | hsa-miR-23b-3p |  |  |  |
|  | hsa-miR-125b-5p |  |  |  |
|  | hsa-miR-141-3p |  |  |  |
|  | hsa-miR-9-5p |  |  |  |
|  | hsa-miR-186-5p |  |  |  |
|  | hsa-miR-155-5p |  |  |  |
|  | hsa-miR-361-5p |  |  |  |
|  | hsa-miR-330-3p |  |  |  |
|  | hsa-miR-429 |  |  |  |

## 4. Discussion

Biomarker studies is crucial for ALS diagnosis, it also shortens the time required to diagnose ALS. *In silico* studies, in particular, play an important role in obtaining a more specific result on biomarkers. In this study, *in silico* methods were used, and nine target genes related to ALS and mitochondrial dysfunction were identified from the literature. A Venn diagram was used to reveal common miRNAs obtained from four different databases for identified genes (Figure 5). We wanted to strengthen our findings by selecting at least three genes targeted for each of the 28 miRNAs. Pathway analyses were performed with the determined genes through Enrichr, g:Profiler and CROssBAR tools. miRNA prediction databases revealed miRNAs that target the identified genes. Running the databases resulted in the prediction of 28 miRNAs. The 28 miRNAs were subjected to miRNA set analyses, and three miRNAs came to the forefront in relation to disease pathogenesis.

In 1993 Bowling and colleagues found that the activity of complex I of the mitochondrial electron transport chain (ETC) increased in brain tissues obtained post-mortem from ALS patients. It was the first report associated with ETC dysfunction in ALS (Bowling et al., 1993). Magrane and colleagues used transgenic mice with SOD1 and TDP-43 mutations, revealed abnormalities in mitochondrial morphology in motor neurons and mitochondrial transport disorders along the axons (Magrané et al., 2014). Researchers investigating where motor neuron death begins have encountered mitochondrial dysfunction. As a result of the studies, it has been reported that there are problems with mitochondrial axonal transport, mitochondria with anomalies accumulated at the neuromuscular junction (NMJ), and motor neuron degeneration associated with them first began from NMJ (Fischer et al., 2004; Magrané et al., 2012; Schaefer et al., 2005). The role and importance of functional disorders occurring in mitochondria in the pathophysiology of ALS have been demonstrated by studies. Researchers trying to unravel the relationship between ALS and mitochondria are investigating which genes are effective in this regard and how they affect it. Considering all these studies *SOD1, FUS, TARDBP, C9orf72, CHCHD10, OPTN, VCP, TBK1* and *BCL2* genes have been associated with mitochondrial dysfunction in ALS (Table 1).

Studies have shown that the mutant SOD1 aggregated in neuron mitochondria causes neuronal degeneration (Vijayvergiya et al., 2005) and clustered mutant SOD1 affects mitochondrial morphology [39], also they aggregate together in the mitochondria of neurons by combining with Bcl-2, which is an antiapoptotic protein (Pasinelli et al., 2004). Also, it has been noted that mutant TDP-43 proteins damage ETC subunit complex I as an aggregate in the neuron mitochondria and cytosol, and they are vulnerable to oxidative stress caused by mutant TDP-43 aggregation (Barmada et al., 2010; Wang et al., 2016; Zuo et al., 2021). Mutant FUS causes functional changes and neurodegeneration in NMJ, and mitochondria enlarged, cristae and membrane structures misaligned also muscle degeneration occurs due to abnormalities in the mitochondria of NMJ and inability to stimulate (Sharma et al., 2016). Studies show that ALS-associated mutant C9orf72 binds to mitochondrial proteins and causes mitochondrial dysfunction, increases oxidative stress, affects mitochondrial membrane potential, decreases Bcl-2 levels, and changes mitochondrial morphology (Dafinca et al., 2016; Lopez-Gonzalez et al., 2016). Furthermore, C9orf72 and BCL2 were found to be significantly associated with autophagy as a result of the pathway enrichment analyses performed in this study.

Mutant OPTN associated with ALS causes mitochondrial dysfunction by accumulating mitochondria in neurons (Wong and Holzbaur, 2014). OPTN is known to supress NF-κB, mutant OPTN cannot supress NF-κB and causes the release of proinflammatory cytokines and neuronal cell death (Akizuki et al., 2013; Liu et al., 2018). TBK1 and OPTN are critical for mitophagy. This information is also supported by the pathway analysis studies conducted here. Mutant OPTN and mutant TBK1 protein also accumulate in the mitochondria, causing ROS and depolarization, mitophagy and autophagosome are damaged (Moore and Holzbaur, 2016). Mutant TANK binding kinase 1 (TBK1) protein creates deficiency for mitophagy, causes accumulation of damaged mitochondria and stress, and affects the pathophysiology of ALS (Harding et al., 2021; Oakes et al., 2017). TBK1 is associated with both autophagy and mitophagy. The pathway analysis studies conducted here also support this information. The mutant CHCHD10 causes mitochondrial fragmentation, mis regulation of cristae structure, and the emergence of different abnormalities (Bannwarth et al., 2014). Many studies revealed that mitochondrial dysfunction caused by mutant CHCHD10 brings about ALS, and death of motor neurons occurs due to the stress (Genin et al., 2019). Valosin-containing protein (VCP) is important for PINK1/Parkin-mediated mitophagy. Mutant VCP associated with ALS damages mitophagy, reducing mitochondria oxygen consumption, lowering the cell’s energy capacity, making the cell vulnerable to processes (Bartolome et al., 2013; Johnson et al., 2010; Kim et al., 2013).

BCL2 key regulator in autophagy, apoptosis, and mitochondrial function dynamics (Akhtar et al., 2004; Zhou et al., 2015). When looking at the BCL2 and motor neuron relationship, later deep motor neuron losses were detected, and it has been shown that this is important for the preservation of specific neuronal populations in the early postnatal period (Michaelidis et al., 1996). The relationship between BCL2 and apoptosis is well known, as is the relationship between apoptosis and ALS. Bcl-2 family members are also known to regulate the mitochondrial metabolic pathway of apoptosis [60–62]. In the literature, BCL2 is not included in the ALS genes and has not been directly associated (https://alsod.ac.uk; Yang et al., 2021; Al-chalabi and Hardiman, 2013) (Nguyen et al., 2018). Changes in the expression levels of the BCL2 and BAX genes were detected in the spinal cord motor neurons of ALS patients in a 1996 study. When compared to controls, BCL2 expression was lower. They also argued that changes in BCL2 levels provide information about the prognosis of ALS disease (Mu et al., 1996). Thus, the levels of miRNA interacting with BCL2 are expected to be differ. Here, in the pathway analyzes, BCL2 is associated with the ALS KEGG pathway and other related pathways (Figure 2). ALS patients’ neurons, an abnormal relationship was detected between SOD1 and BCL2. It was revealed that they together undergo precipitation in mitochondria, and decrease mitochondria ADP permeability (Pasinelli et al., 2004; Pedrini et al., 2010; Tan et al., 2013). In pathway analysis, the autophagy, apoptosis, p53 signalling, and neurodegeneration pathways were discovered to be significantly related (Figure 3, Table 2). Here, genes that affect mitochondrial dysfunction are gathered under a single heading. The miRNAs that we identified targeting all these gene products are also found together in the present study.

**Figure 2:**
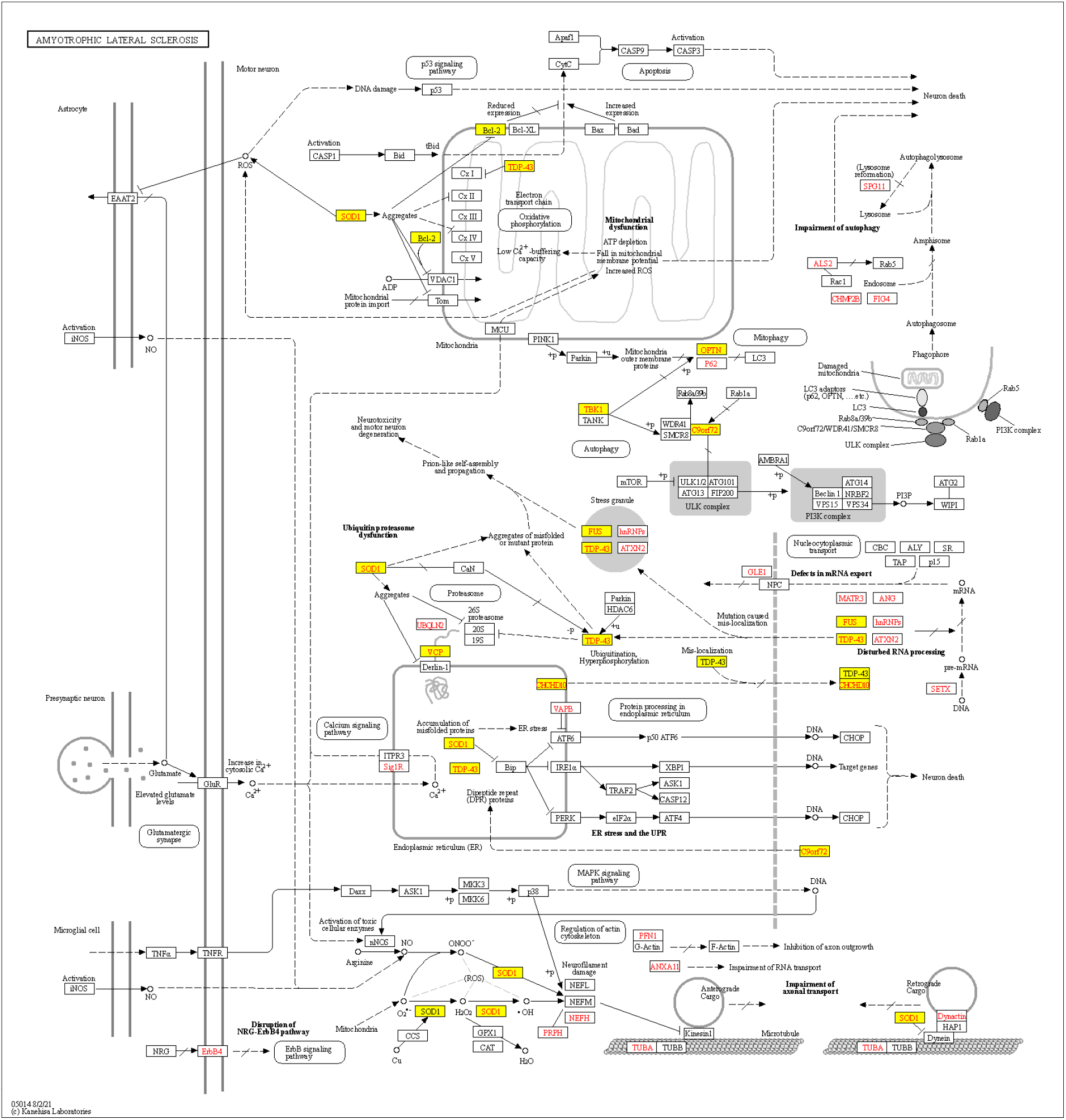
KEGG, Amyotrophic Lateral Sclerosis Pathway, Map05014. The yellow marked nodes represent genes which are related to mitochondrial dysfunction in ALS.

**Figure 3:**
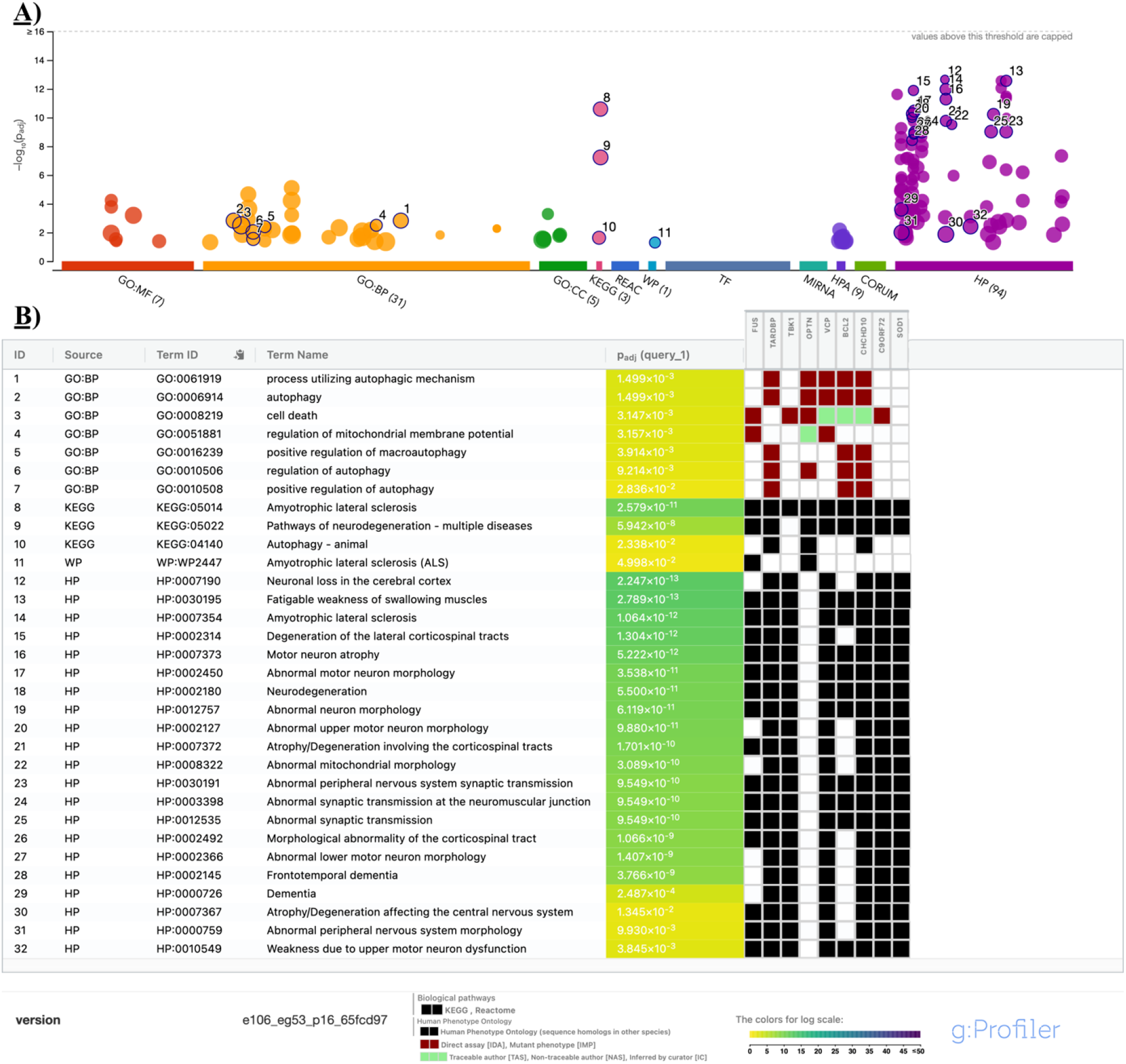
Pathway Enrichment Analyses with tools. A) Pathway enrichment analysis for using g: Profiler tool. B) Detailed pathway enrichment analysis with the genes.

**Figure 4:**
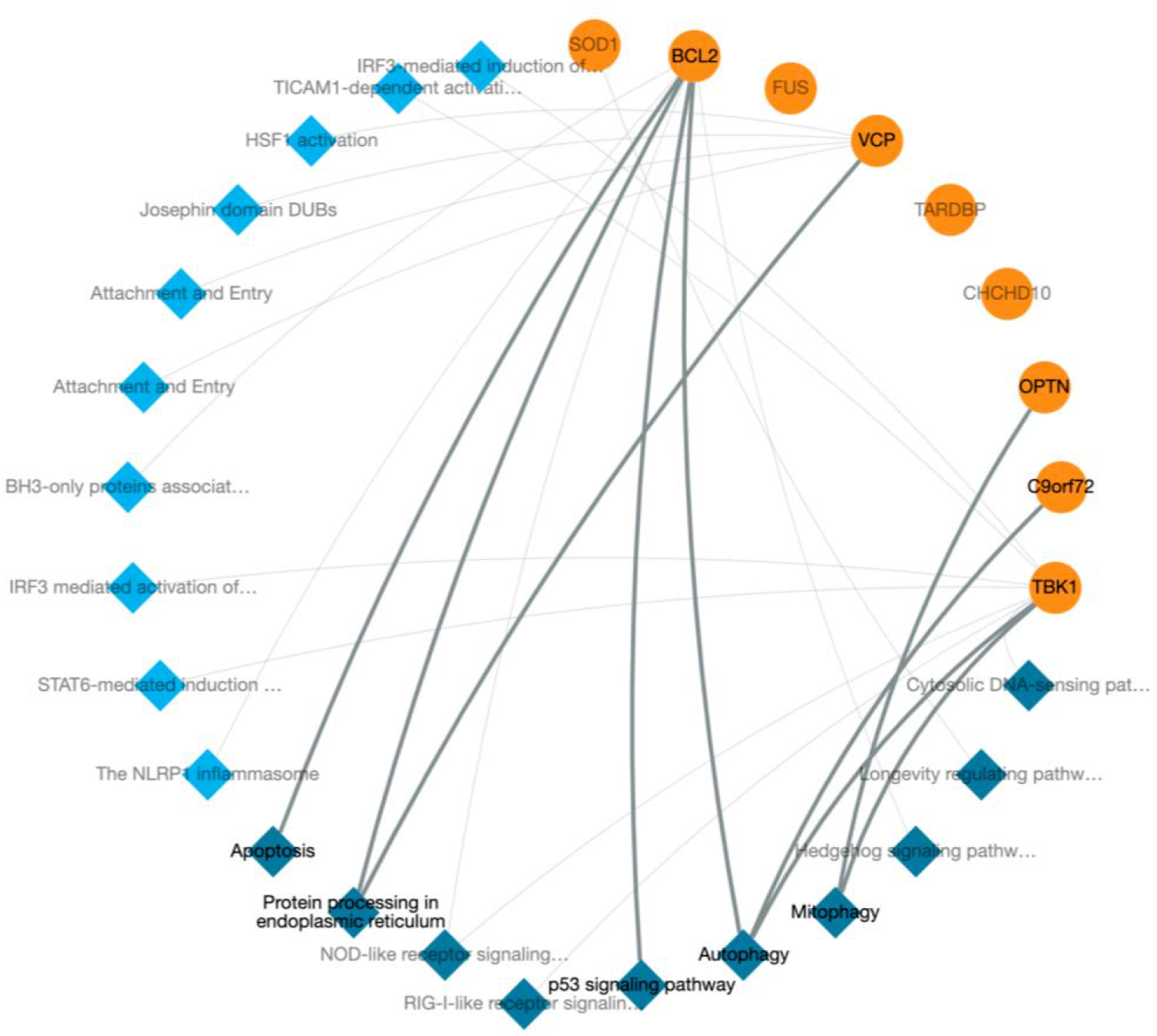
Pathway Enrichment Analysis with CROssBAR tool.

**Figure 5:**
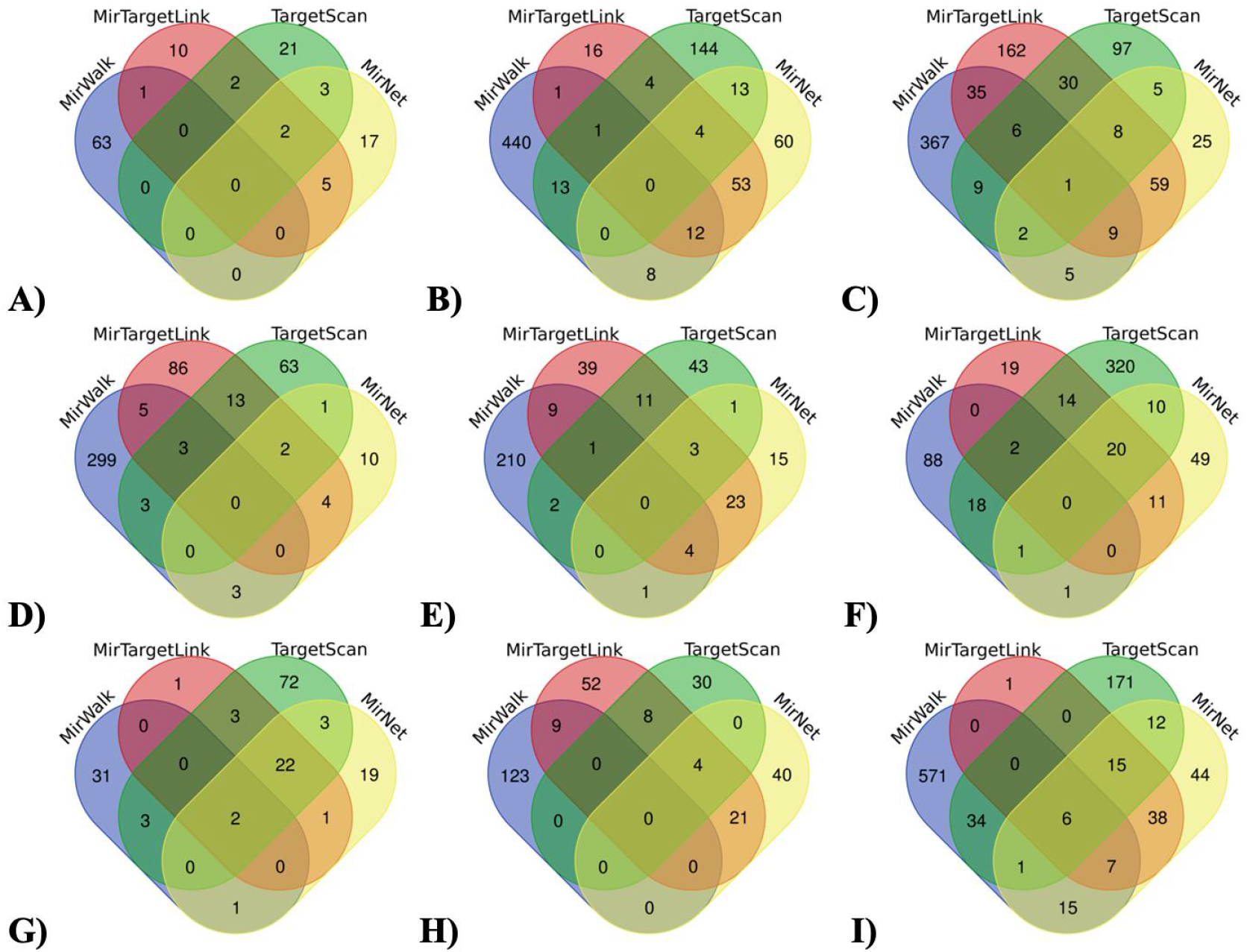
Venn diagram images used to reveal the common miRNAs obtained using four different databases for the identified genes. There are 6 miRNAs for *BCL2* common to all for databases. There is 1 miRNA common to the four databases for *TARDBP*. There are 2 common miRNAs for *CHCHD10* in all four databases. There are common miRNAs in max 3 databases for the remaining *SOD1, FUS, C9ORF72, TBK1, OPTN, VCP* genes. There are no common miRNAs for these 6 genes in the four databases. A: SOD1, B: FUS, C: TARDBP, D: C9ORF72, E: OPTN, F: VCP, G: CHCHD10, H: TBK1, I: BCL2

miR-9 is highly abundant in the brain region and is a brain-specific miRNA. It is also reported that higher level of mir-9 in brain affects especially development in the embryonic period compared to the adult periods (Landgraf et al., 2007; Miska et al., 2004; Sempere et al., 2004). Furthermore, it has an important role in neurogenesis and neuronal differentiation (Leucht et al., 2008). miR-9 enables neural stem cells to self-renewable and regulates the proliferation of neural progenitor cells (Shibata et al., 2011, 2008; Zhao et al., 2009). According to the miRNA set analyses, miR-9 is significantly associated with the brain, neuron differentiation, and brain development, which is consistent with this information (Table 3). Researchers, who think that changes in the mir-9 expression level found to be important for neural development and differentiation, may cause neurodegeneration. miR-9 has been suggested to be a potential regulator of mature neurons and neurodegeneration (Haramati et al., 2010). In studies with various cell cultures and model organisms associated with ALS, it has been determined that miR-9 is associated with neurodegeneration and changes in its expression levels (Marcuzzo et al., 2015; Zhang et al., 2013; Zhou et al., 2013). Therefore, miR-9 may be a potential biomarker for ALS and must be further investigated

The miRNA-200 family, of which miR-141 is a member, is found in two different genomic regions with its five members (miR-200a, miR-200b, miR-200c, miR-429 and miR-141) (Huang et al., 2014). miR-141 has been shown to modulate apoptosis, and its expression is increased after fibroblasts are exposed to oxidative stress (Mateescu et al., 2011). miR-141, a ROS-induced miRNA, has been found to be overexpressed in the brains of patients with Alzheimer’s disease (Cogswell et al., 2008). A study confirmed that miR-141 and miR-9 expression is significantly increased in Parkinson’s disease in response to oxidative stress (Rostamian Delavar et al., 2018). Furthermore, in another study, they discovered that miR-9 and miR-141 could be potential biomarkers in acute pancreatitis patients’ serum data (Lu et al., 2017). Here, we have summarized the roles of miR-141.

miR-125b is microglia enriched miRNA (Butovsky et al., 2014; Parisi et al., 2016) and miR-125b is one of the neuronal miRNAs (Lee et al., 2005). Furthermore, two ALS causative genes, TDP43 and FUS, are specifically involved in miR-125b processing, which contributes to the association of this specific miRNA with ALS disease (di Carlo et al., 2013; Morlando et al., 2012). Our findings are in line with previous research performed on showing that miR-125b targets the TARDBP and FUS genes (see Supplementary Table 1). In mice with the ALS SOD1 mutation, miR-125b was found to be up-regulated in microglia cells (Parisi et al., 2013). In contrast to Parisi et al., miR-125b was found to be down-regulated in ALS patients (Saucier et al., 2019). It was also discovered to be downregulated in the NMJ of sALS patients (de Felice et al., 2018). In the study of Giroud et al., in vitro, miR-125b-5p overexpression and knockdown decreased and increased mitochondrial biogenesis, in both. According to these findings, miR-125b influences mitochondrial biogenesis and appears to play an important role in the regulation of mitochondrial biogenesis (Giroud et al., 2016). Also, in another study performed that miR-125b reduces mitochondrial respiration (Duroux-Richard et al., 2016). Bcl-2 is targeted by miR-125b (Ouyang and Giffard, 2014; Shi et al., 2012). Our findings are in line with previous research performed on showing that miR-125b targets BCL2 gene (see Supplementary Table 1). These results strengthen our results in supplementary table 1. All these findings strengthen the role of miR-9, miR-141, and miR-125b miRNAs in ALS pathology.

We argue that the levels of expression of these miRNAs may differ significantly in neurodegeneration. It is widely known that research is being conducted on the use of miRNAs as biomarkers to shorten and facilitate the diagnosis of many diseases. The average duration of diagnosis of ALS disease is 12 months. In relation to this situation, biomarker research has priority in ALS studies (Joilin et al., 2020; Takahashi et al., 2015; Toivonen et al., 2014).

## 5. Conclusions

Overall, we demonstrated that the genes involved in mitochondrial dysfunction in ALS based on the literature. Here, we showed predicted effects of miR-9, and miR-141 targeting the identified genes on ALS by *in silico* methods. Further studies are required to investigate and confirm the identified miRNAs and their target genes for clinical use as biomarkers.

## Author Contributions

Baykal, Gulcin carried out methodology, data curation, writing, drafted the manuscript, conceptualization. Erkal, Burcin carried out writing, editing, revised the manuscript, conceptualization. Vural Korkut, Senay carried out writing, editing, revised the manuscript, conceptualization. All authors read and approved the final manuscript.

## Supplementary Materials

**supplement Table S1:**
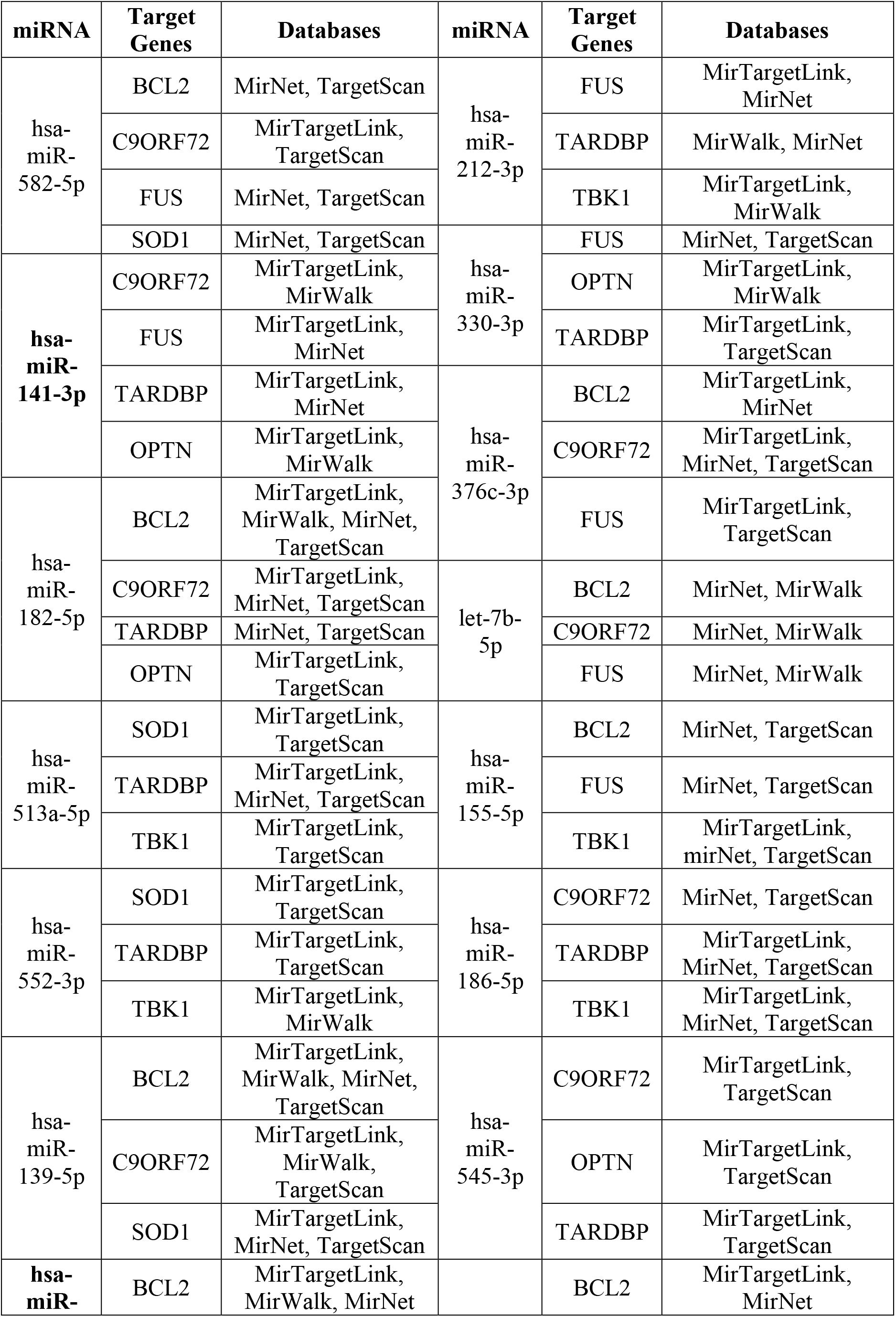
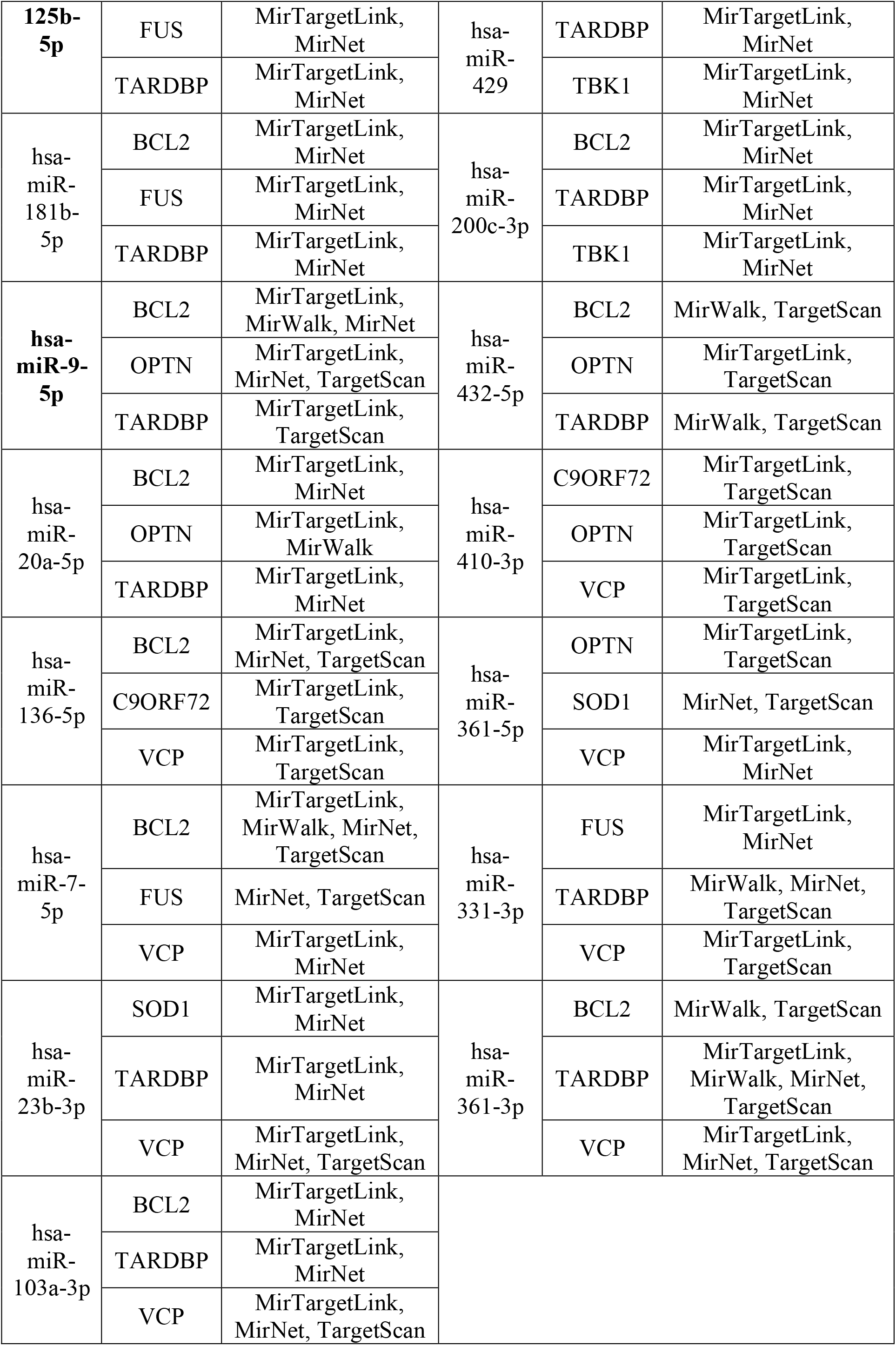
miRNAs Targeting Genes Associated with Mitochondrial Dysfunction

## References

Åkerblom, M., Sachdeva, R., Barde, I., Verp, S., Gentner, B., Trono, D., Jakobsson, J., 2012. Journal of Neuroscience 32, 8879–8889.

Akhtar, R.S., Ness, J.M., Roth, K.A., 2004. Bcl-2 family regulation of neuronal development and neurodegeneration. Biochimica et Biophysica Acta - Molecular Cell Research.

Akizuki, M., Yamashita, H., Uemura, K., Maruyama, H., Kawakami, H., Ito, H., Takahashi, R., 2013. Journal of Neurochemistry 126, 699–704.

Atsumi, T., 1981. Acta Neuropathologica 55, 193–198.

Bannwarth, S., Ait-El-Mkadem, S., Chaussenot, A., Genin, E.C., Lacas-Gervais, S., Fragaki, K., Berg-Alonso, L., Kageyama, Y., Serre, V., Moore, D.G., Verschueren, A., Rouzier, C., le Ber, I., Augé, G., Cochaud, C., Lespinasse, F., N’guyen, K., de Septenville, A., Brice, A., Yu-Wai-Man, P., Sesaki, H., Pouget, J., Paquis-Flucklinger, V., 2014. Brain 137, 2329–2345.

Barmada, S.J., Skibinski, G., Korb, E., Rao, E.J., Wu, J.Y., Finkbeiner, S., 2010. Journal of Neuroscience 30, 639–649.

Bartel, D.P., 2004. Review MicroRNAs: Genomics, Biogenesis, Mechanism, and Function ulation of hematopoietic lineage differentiation in mam-mals (Chen et al., 2004), and control of leaf and flower development in plants (Aukerman and Sakai, 2003, Cell.

Bartel, D.P., 2009. Cell 136, 215–233.

Bartolome, F., Wu, H.C., Burchell, V.S., Preza, E., Wray, S., Mahoney, C.J., Fox, N.C., Calvo, A., Canosa, A., Moglia, C., Mandrioli, J., Chiò, A., Orrell, R.W., Houlden, H., Hardy, J., Abramov, A.Y., Plun-Favreau, H., 2013. Neuron 78, 57–64.

Benjaminsen, E., Alstadhaug, K.B., Gulsvik, M., Baloch, F.K., Odeh, F., 2018. Amyotrophic Lateral Sclerosis and Frontotemporal Degeneration 19, 522–527.

Bordoni, L., Gabbianelli, R., 2020. Mitochondrial dna and neurodegeneration: Any role for dietary antioxidants? Antioxidants.

Bowling, A.C., Schulz, J.B., Brown, R.H., Beal, M.F., 1993. Journal of Neurochemistry 61, 2322–2325.

Bowser, R., Turner, M.R., Shefner, J., 2011. Biomarkers in amyotrophic lateral sclerosis: Opportunities and limitations. Nature Reviews Neurology.

Brown, R.H., Al-Chalabi, A., 2017. New England Journal of Medicine 377, 162–172.

Butovsky, O., Jedrychowski, M.P., Moore, C.S., Cialic, R., Lanser, A.J., Gabriely, G., Koeglsperger, T., Dake, B., Wu, P.M., Doykan, C.E., Fanek, Z., Liu, L., Chen, Z., Rothstein, J.D., Ransohoff, R.M., Gygi, S.P., Antel, J.P., Weiner, H.L., 2014. Nature Neuroscience 17, 131–143.

di Carlo, V., Grossi, E., Laneve, P., Morlando, M., Dini Modigliani, S., Ballarino, M., Bozzoni, I., Caffarelli, E., 2013. TDP-43 regulates the microprocessor complex activity during in vitro neuronal differentiation. Molecular Neurobiology.

Cogswell, J.P., Ward, J., Taylor, I.A., Waters, M., Shi, Y., Cannon, B., Kelnar, K., Kemppainen, J., Brown, D., Chen, C., Prinjha, R.K., Richardson, J.C., Saunders, A.M., Roses, A.D., Richards, C.A., 2008. Journal of Alzheimer’s Disease 14, 27–41.

Conaco, C., Otto, S., Han, J.-J., Mandel, G., 2006. Reciprocal actions of REST and a microRNA promote neuronal identity.

Dafinca, R., Scaber, J., Ababneh, N., Lalic, T., Weir, G., Christian, H., Vowles, J., Douglas, A.G.L., Fletcher-Jones, A., Browne, C., Nakanishi, M., Turner, M.R., Wade-Martins, R., Cowley, S.A., Talbot, K., 2016. Stem Cells 34, 2063–2078.

Duroux-Richard, I., Roubert, C., Ammari, M., PrésumeyPr, J., GrünGr, J.R., GrützkauGr, A., Lecellier, C.-H., erie Boitez, V., Codogno, P., Escoubet, J., Pers, Y.-M., Jorgensen, C., Apparailly, F., 2016.

van Es, M.A., Hardiman, O., Chio, A., Al-Chalabi, A., Pasterkamp, R.J., Veldink, J.H., van den Berg, L.H., 2017. The Lancet 390, 2084–2098.

de Felice, B., Manfellotto, F., Fiorentino, G., Annunziata, A., Biffali, E., Pannone, R., Federico, A., 2018. Frontiers in Genetics 9.

Fischer, L.R., Culver, D.G., Tennant, P., Davis, A.A., Wang, M., Castellano-Sanchez, A., Khan, J., Polak, M.A., Glass, J.D., 2004. Experimental Neurology 185, 232–240.

Genin, E.C., Madji Hounoum, B., Bannwarth, S., Fragaki, K., Lacas-Gervais, S., Mauri-Crouzet, A., Lespinasse, F., Neveu, J., Ropert, B., Augé, G., Cochaud, C., Lefebvre-Omar, C., Bigou, S., Chiot, A., Mochel, F., Boillée, S., Lobsiger, C.S., Bohl, D., Ricci, J.E., Paquis-Flucklinger, V., 2019. Acta Neuropathologica.

Giroud, M., Pisani, D.F., Karbiener, M., Barquissau, V., Ghandour, R.A., Tews, D., Fischer-Posovszky, P., Chambard, J.C., Knippschild, U., Niemi, T., Taittonen, M., Nuutila, P., Wabitsch, M., Herzig, S., Virtanen, K.A., Langin, D., Scheideler, M., Amri, E.Z., 2016. Molecular Metabolism 5, 615–625.

Grimson, A., Farh, K.K.H., Johnston, W.K., Garrett-Engele, P., Lim, L.P., Bartel, D.P., 2007. Molecular Cell 27, 91–105.

Haramati, S., Chapnik, E., Sztainberg, Y., Eilam, R., Zwang, R., Gershoni, N., McGlinn, E., Heiser, P.W., Wills, A.M., Wirguin, I., Rubin, L.L., Misawa, H., Tabin, C.J., Brown, R., Chen, A., Hornstein, E., 2010. Proc Natl Acad Sci U S A 107, 13111–13116.

Hardiman, O., Al-Chalabi, A., Chio, A., Corr, E.M., Logroscino, G., Robberecht, W., Shaw, P.J., Simmons, Z., van den Berg, L.H., 2017. Nature Reviews Disease Primers 3, 17071.

Harding, O., Evans, C.S., Ye, J., Cheung, J., Maniatis, T., Holzbaur, E.L.F., 2021. Proceedings of the National Academy of Sciences 118, e2025053118.

Huang, H.N., Chen, S.Y., Hwang, S.M., Yu, C.C., Su, M.W., Mai, W., Wang, H.W., Cheng, W.C., Schuyler, S.C., Ma, N., Lu, F.L., Lu, J., 2014. Stem Cell Research 12, 338–353.

Jaiswal, M.K., 2017. Frontiers in Cellular Neuroscience 10.

Johnson, J.O., Mandrioli, J., Benatar, M., Abramzon, Y., van Deerlin, V.M., Trojanowski, J.Q., Gibbs, J.R., Brunetti, M., Gronka, S., Wuu, J., Ding, J., McCluskey, L., Martinez-Lage, M., Falcone, D., Hernandez, D.G., Arepalli, S., Chong, S., Schymick, J.C., Rothstein, J., Landi, F., Wang, Y.D., Calvo, A., Mora, G., Sabatelli, M., Monsurrò, M.R., Battistini, S., Salvi, F., Spataro, R., Sola, P., Borghero, G., Galassi, G., Scholz, S.W., Taylor, J.P., Restagno, G., Chiò, A., Traynor, B.J., 2010. Neuron 68, 857–864.

Joilin, G., Gray, E., Thompson, A.G., Bobeva, Y., Talbot, K., Weishaupt, J., Ludolph, A., Malaspina, A., Leigh, P.N., Newbury, S.F., Turner, M.R., Hafezparast, M., 2020. Brain Communications 2.

Jun, K.Y., Park, J., Oh, K.-W., Kim, E.M., Bae, J.S., Kim, I., Kim, S.H., 2019. Journal of Neurology, Neurosurgery & Psychiatry 90, 395–403.

Kim, N.C., Tresse, E., Kolaitis, R.M., Molliex, A., Thomas, R.E., Alami, N.H., Wang, B., Joshi, A., Smith, R.B., Ritson, G.P., Winborn, B.J., Moore, J., Lee, J.Y., Yao, T.P., Pallanck, L., Kundu, M., Taylor, J.P., 2013. Neuron 78, 65–80.

Landgraf, P., Rusu, M., Sheridan, R., Sewer, A., Iovino, N., Aravin, A., Pfeffer, S., Rice, A., Kamphorst, A.O., Landthaler, M., Lin, C., Socci, N.D., Hermida, L., Fulci, V., Chiaretti, S., Foà, R., Schliwka, J., Fuchs, U., Novosel, A., Müller, R.-U., Schermer, B., Bissels, U., Inman, J., Phan, Q., Chien, M., Weir, D.B., Choksi, R., de Vita, G., Frezzetti, D., Trompeter, H.-I., Hornung, V., Teng, G., Hartmann, G., Palkovits, M., di Lauro, R., Wernet, P., Macino, G., Rogler, C.E., Nagle, J.W., Ju, J., Papavasiliou, F.N., Benzing, T., Lichter, P., Tam, W., Brownstein, M.J., Bosio, A., Borkhardt, A., Russo, J.J., Sander, C., Zavolan, M., Tuschl, T., 2007. Cell 129, 1401–1414.

Lee, Y.S., Kim, H.K., Chung, S., Kim, K.S., Dutta, A., 2005. Journal of Biological Chemistry 280, 16635–16641.

Leighton, D.J., Newton, J., Stephenson, L.J., Colville, S., Davenport, R., Gorrie, G., Morrison, I., Swingler, R., Chandran, S., Pal, S., 2019. Journal of Neurology 266, 817–825.

Leucht, C., Stigloher, C., Wizenmann, A., Klafke, R., Folchert, A., Bally-Cuif, L., 2008. Nature Neuroscience 11, 641–648.

Lezi, E., Swerdlow, R.H., 2012. Advances in Experimental Medicine and Biology 942, 269–286.

Lin, J., Diamanduros, A., Chowdhury, S.A., Scelsa, S., Latov, N., Sadiq, S.A., 2009. Journal of Neurology 256, 774–782.

Liu, Z., Li, H., Hong, C., Chen, M., Yue, T., Chen, C., Wang, Z., You, Q., Li, C., Weng, Q., Xie, H., Hu, R., 2018. Frontiers in Immunology 9.

Longinetti, E., Regodón Wallin, A., Samuelsson, K., Press, R., Zachau, A., Ronnevi, L.-O., Kierkegaard, M., Andersen, P.M., Hillert, J., Fang, F., Ingre, C., 2018. Amyotrophic Lateral Sclerosis and Frontotemporal Degeneration 19, 528–537.

Lopez-Gonzalez, R., Lu, Y., Gendron, T.F., Karydas, A., Tran, H., Yang, D., Petrucelli, L., Miller, B.L., Almeida, S., Gao, F.B., 2016. Neuron 92, 383–391.

Lu, P., Wang, F., Wu, J., Wang, C., Yan, J., Li, Z., Song, J., Wang, J., 2017. Disease Markers 2017, 1–8.

Magrané, J., Cortez, C., Gan, W.B., Manfredi, G., 2014. Human Molecular Genetics 23, 1413–1424.

Magrané, J., Sahawneh, M.A., Przedborski, S., Estévez, Á.G., Manfredi, G., 2012. Journal of Neuroscience 32, 229–242.

Marcuzzo, S., Bonanno, S., Kapetis, D., Barzago, C., Cavalcante, P., D’Alessandro, S., Mantegazza, R., Bernasconi, P., 2015. Molecular Brain 8.

Mateescu, B., Batista, L., Cardon, M., Gruosso, T., de Feraudy, Y., Mariani, O., Nicolas, A., Meyniel, J.-P., Cottu, P., Sastre-Garau, X., Mechta-Grigoriou, F., 2011. Nature Medicine 17, 1627–1635.

Mejzini, R., Flynn, L.L., Pitout, I.L., Fletcher, S., Wilton, S.D., Akkari, P.A., 2019. Frontiers in Neuroscience 13.

Michaelidis, T.M., Sendtner, M., Cooper, J.D., 1996. Inactivation of bcl-2 Results in Progressive Degeneration of Motoneurons, Sympathetic and Sensory Neurons during Early Postnatal Development, Neuron.

Miska, E.A., Alvarez-Saavedra, E., Townsend, M., Yoshii, A., Šestan, N., Rakic, P., Constantine-Paton, M., Horvitz, R., 2004. Open Access Microarray analysis of microRNA expression in the developing mammalian brain.

Moore, A.S., Holzbaur, E.L.F., 2016. Proc Natl Acad Sci U S A 113, E3349–E3358.

Morgan, S., Orrell, R.W., 2016. British Medical Bulletin 119, 87–98.

Morlando, M., Dini Modigliani, S., Torrelli, G., Rosa, A., di Carlo, V., Caffarelli, E., Bozzoni, I., 2012. EMBO Journal 31, 4502–4510.

Mu, X., He, J., Anderson, D.W., Trojanowski, J.Q., Springer, J.E., 1996. Annals of Neurology 40, 379–386.

Nguyen, H.P., van Broeckhoven, C., van der Zee, J., 2018. ALS Genes in the Genomic Era and their Implications for FTD. Trends in Genetics.

Oakes, J.A., Davies, M.C., Collins, M.O., 2017. Molecular Brain 10, 1–10.

Ola, M.S., Nawaz, M., Ahsan, H., 2011. Role of Bcl-2 family proteins and caspases in the regulation of apoptosis. Molecular and Cellular Biochemistry.

Ouyang, Y.B., Giffard, R.G., 2014. MicroRNAs affect BCL-2 family proteins in the setting of cerebral ischemia. Neurochemistry International.

Palese, F., Sartori, A., Verriello, L., Ros, S., Passadore, P., Manganotti, P., Barbone, F., Pisa, F.E., 2019. Amyotrophic Lateral Sclerosis and Frontotemporal Degeneration 20, 90–99.

Parisi, C., Arisi, I., D’Ambrosi, N., Storti, A.E., Brandi, R., D’Onofrio, M., Volonté, C., 2013. Cell Death and Disease 4.

Parisi, C., Napoli, G., Amadio, S., Spalloni, A., Apolloni, S., Longone, P., Volonte, C., 2016. Cell Death and Differentiation 23, 531–541.

Pasinelli, P., Belford, M.E., Lennon, N., Bacskai, B.J., Hyman, B.T., Trotti, D., Brown, R.H., 2004. Neuron 43, 19–30.

Pedrini, S., Sau, D., Guareschi, S., Bogush, M., Brown, R.H., Naniche, N., Kia, A., Trotti, D., Pasinelli, P., 2010. Human Molecular Genetics 19, 2974–2986.

Peoples, J.N., Saraf, A., Ghazal, N., Pham, T.T., Kwong, J.Q., 2019. Experimental & Molecular Medicine 51, 1–13.

Peter, M.E., 2010. Oncogene 29, 2161–2164.

Rose, L., McKim, D., Leasa, D., Nonoyama, M., Tandon, A., Bai, Y.Q., Amin, R., Katz, S., Goldstein, R., Gershon, A., 2019. PLOS ONE 14, e0210574.

Rostamian Delavar, M., Baghi, M., Safaeinejad, Z., Kiani-Esfahani, A., Ghaedi, K., Nasr-Esfahani, M.H., 2018. Gene 662, 54–65.

Sasaki, S., Iwata, M., 1996. Neuroscience Letters 204, 53–56.

Sasaki, S., Iwata, M., 2007. Journal of Neuropathology and Experimental Neurology 66, 10–16.

Sathasivam, S., Ince, P.G., Shaw, P.J., 2001. Apoptosis in amyotrophic lateral sclerosis: A review of the evidence. Neuropathology and Applied Neurobiology.

Saucier, D., Wajnberg, G., Roy, J., Beauregard, A.P., Chacko, S., Crapoulet, N., Fournier, S., Ghosh, A., Lewis, S.M., Marrero, A., O’Connell, C., Ouellette, R.J., Morin, P., 2019. Brain Research 1708, 100–108.

Schaefer, A.M., Sanes, J.R., Lichtman, J.W., 2005. The Journal of Comparative Neurology 490, 209–219.

Sempere, L.F., Freemantle, S., Pitha-Rowe, I., Moss, E., Dmitrovsky, E., Ambros, V., 2004. Expression profiling of mammalian microRNAs uncovers a subset of brain-expressed microRNAs with possible roles in murine and human neuronal differentiation, Sempere et al.

Sharma, A., Lyashchenko, A.K., Lu, L., Nasrabady, S.E., Elmaleh, M., Mendelsohn, M., Nemes, A., Tapia, J.C., Mentis, G.Z., Shneider, N.A., 2016. Nature Communications 7.

Shi, L., Zhang, S., Feng, K., Wu, F., Wan, Y., Wang, Z., Zhang, J., Wang, Y., Yan, W., Fu, Z., You, Y., 2012. International Journal of Oncology 40, 119–129.

Shibata, M., Kurokawa, D., Nakao, H., Ohmura, T., Aizawa, S., 2008. Journal of Neuroscience 28, 10415–10421.

Shibata, M., Nakao, H., Kiyonari, H., Abe, T., Aizawa, S., 2011. Journal of Neuroscience 31, 3407–3422.

Tadic, V., Prell, T., Lautenschlaeger, J., Grosskreutz, J., 2014. Frontiers in Cellular Neuroscience 8.

Takahashi, I., Hama, Y., Matsushima, M., Hirotani, M., Kano, T., Hohzen, H., Yabe, I., Utsumi, J., Sasaki, H., 2015. Molecular Brain 8, 67.

Tan, W., Naniche, N., Bogush, A., Pedrini, S., Trotti, D., Pasinelli, P., 2013. Journal of Neuroscience 33, 11588–11598.

Toivonen, J.M., Manzano, R., Oliván, S., Zaragoza, P., García-Redondo, A., Osta, R., 2014. PLoS ONE 9, e89065.

Troost, D., Aten, J., Morsink, F., de Jong, J.M.B.V., 1995. Neuropathology and Applied Neurobiology 21, 498–504.

vande Velde, C., McDonald, K.K., Boukhedimi, Y., McAlonis-Downes, M., Lobsiger, C.S., Hadj, S.B., Zandona, A., Julien, J.P., Shah, S.B., Cleveland, D.W., 2011. PLoS ONE 6.

Vijayvergiya, C., Beal, M.F., Buck, J., Manfredi, G., 2005. Journal of Neuroscience 25, 2463–2470.

Wallace, D.C., 2012. Nature Reviews Cancer 12, 685–698.

Wang, W., Wang, L., Lu, J., Siedlak, S.L., Fujioka, H., Liang, J., Jiang, S., Ma, X., Jiang, Z., da Rocha, E.L., Sheng, M., Choi, H., Lerou, P.H., Li, H., Wang, X., 2016. Nature Medicine 22, 869–878.

Wong, Y.C., Holzbaur, E.L.F., 2014. Proc Natl Acad Sci U S A 111, E4439–E4448.

Wu, S., Huang, S., Ding, J., Zhao, Y., Liang, L., Liu, T., Zhan, R., He, X., 2010. Oncogene 29, 2302–2308.

Zhang, Z., Almeida, S., Lu, Y., Nishimura, A.L., Peng, L., Sun, D., Wu, B., Karydas, A.M., Tartaglia, M.C., Fong, J.C., Miller, B.L., Farese, R. v., Moore, M.J., Shaw, C.E., Gao, F.B., 2013. PLoS ONE 8.

Zhao, C., Sun, G., Li, S., Shi, Y., 2009. Nature Structural & Molecular Biology 16, 365–371.

Zhou, F., Guan, Y., Chen, Y., Zhang, C., Yu, L., Gao, H., Du, H., Liu, B., Wang, X., 2013. Int J Clin Exp Pathol 6, 1826–38.

Zhou, L., Wang, H., Ren, H., Hu, Q., Ying, Z., Wang, G., 2015. Molecular Neurobiology 52, 1180–1189.

Zhou, S., Zhou, Y., Qian, S., Chang, W., Wang, L., Fan, D., 2018. Brain and Behavior 8, e01131.

Zuo, X., Zhou, J., Li, Y., Wu, K., Chen, Z., Luo, Z., Zhang, X., Liang, Y., Esteban, M.A., Zhou, Y., Fu, X.D., 2021. Nature Structural and Molecular Biology 28, 132–142.

