## Supplemental Table 1 for "*In silico* identification of miRNAs related to mitochondrial dysfunction in amyotrophic lateral sclerosis"

**Table S1:** miRNAs Targeting Genes Associated with Mitochondrial Dysfunction

| miRNA | Target Genes | Databases | miRNA | Target Genes | Databases |
|---|---|---|---|---|---|
| hsa-miR-582-5p | BCL2 | MirNet, TargetScan | hsa-miR-212-3p | FUS | MirTargetLink, MirNet |
| | C9ORF72 | MirTargetLink, TargetScan | | TARDBP | MirWalk, MirNet |
| | FUS | MirNet, TargetScan | | TBK1 | MirTargetLink, MirWalk |
| | SOD1 | MirNet, TargetScan | hsa-miR-330-3p | FUS | MirNet, TargetScan |
| **hsa-miR-141-3p** | C9ORF72 | MirTargetLink, MirWalk | | OPTN | MirTargetLink, MirWalk |
| | FUS | MirTargetLink, MirNet | | TARDBP | MirTargetLink, TargetScan |
| | TARDBP | MirTargetLink, MirNet | hsa-miR-376c-3p | BCL2 | MirTargetLink, MirNet |
| | OPTN | MirTargetLink, MirWalk | | C9ORF72 | MirTargetLink, MirNet, TargetScan |
| hsa-miR-182-5p | BCL2 | MirTargetLink, MirWalk, MirNet, TargetScan | | FUS | MirTargetLink, TargetScan |
| | C9ORF72 | MirTargetLink, MirNet, TargetScan | let-7b-5p | BCL2 | MirNet, MirWalk |
| | TARDBP | MirNet, TargetScan | | C9ORF72 | MirNet, MirWalk |
| | OPTN | MirTargetLink, TargetScan | | FUS | MirNet, MirWalk |
| hsa-miR-513a-5p | SOD1 | MirTargetLink, TargetScan | hsa-miR-155-5p | BCL2 | MirNet, TargetScan |
| | TARDBP | MirTargetLink, MirNet, TargetScan | | FUS | MirNet, TargetScan |
| | TBK1 | MirTargetLink, TargetScan | | TBK1 | MirTargetLink, mirNet, TargetScan |
| hsa-miR-552-3p | SOD1 | MirTargetLink, TargetScan | hsa-miR-186-5p | C9ORF72 | MirNet, TargetScan |
| | TARDBP | MirTargetLink, TargetScan | | TARDBP | MirTargetLink, MirNet, TargetScan |
| | TBK1 | MirTargetLink, MirWalk | | TBK1 | MirTargetLink, MirNet, TargetScan |
| hsa-miR-139-5p | BCL2 | MirTargetLink, MirWalk, MirNet, TargetScan | hsa-miR-545-3p | C9ORF72 | MirTargetLink, TargetScan |
| | C9ORF72 | MirTargetLink, MirWalk, TargetScan | | OPTN | MirTargetLink, TargetScan |
| | SOD1 | MirTargetLink, MirNet, TargetScan | | TARDBP | MirTargetLink, TargetScan |
| **hsa-miR-** | BCL2 | MirTargetLink, MirWalk, MirNet | | BCL2 | MirTargetLink, MirNet |

| | | | | | |
|---|---|---|---|---|---|
| **125b-5p** | FUS | MirTargetLink, MirNet | hsa-miR-429 | TARDBP | MirTargetLink, MirNet |
| | TARDBP | MirTargetLink, MirNet | | TBK1 | MirTargetLink, MirNet |
| hsa-miR-181b-5p | BCL2 | MirTargetLink, MirNet | hsa-miR-200c-3p | BCL2 | MirTargetLink, MirNet |
| | FUS | MirTargetLink, MirNet | | TARDBP | MirTargetLink, MirNet |
| | TARDBP | MirTargetLink, MirNet | | TBK1 | MirTargetLink, MirNet |
| **hsa-miR-9-5p** | BCL2 | MirTargetLink, MirWalk, MirNet | hsa-miR-432-5p | BCL2 | MirWalk, TargetScan |
| | OPTN | MirTargetLink, MirNet, TargetScan | | OPTN | MirTargetLink, TargetScan |
| | TARDBP | MirTargetLink, TargetScan | | TARDBP | MirWalk, TargetScan |
| hsa-miR-20a-5p | BCL2 | MirTargetLink, MirNet | hsa-miR-410-3p | C9ORF72 | MirTargetLink, TargetScan |
| | OPTN | MirTargetLink, MirWalk | | OPTN | MirTargetLink, TargetScan |
| | TARDBP | MirTargetLink, MirNet | | VCP | MirTargetLink, TargetScan |
| hsa-miR-136-5p | BCL2 | MirTargetLink, MirNet, TargetScan | hsa-miR-361-5p | OPTN | MirTargetLink, TargetScan |
| | C9ORF72 | MirTargetLink, TargetScan | | SOD1 | MirNet, TargetScan |
| | VCP | MirTargetLink, TargetScan | | VCP | MirTargetLink, MirNet |
| hsa-miR-7-5p | BCL2 | MirTargetLink, MirWalk, MirNet, TargetScan | hsa-miR-331-3p | FUS | MirTargetLink, MirNet |
| | FUS | MirNet, TargetScan | | TARDBP | MirWalk, MirNet, TargetScan |
| | VCP | MirTargetLink, MirNet | | VCP | MirTargetLink, TargetScan |
| hsa-miR-23b-3p | SOD1 | MirTargetLink, MirNet | hsa-miR-361-3p | BCL2 | MirWalk, TargetScan |
| | TARDBP | MirTargetLink, MirNet | | TARDBP | MirTargetLink, MirWalk, MirNet, TargetScan |
| | VCP | MirTargetLink, MirNet, TargetScan | | VCP | MirTargetLink, MirNet, TargetScan |
| hsa-miR-103a-3p | BCL2 | MirTargetLink, MirNet | | | |
| | TARDBP | MirTargetLink, MirNet | | | |
| | VCP | MirTargetLink, MirNet, TargetScan | | | |
